# Uncovering organizing principles of protein-protein interaction networks

**DOI:** 10.1101/2024.03.20.585897

**Authors:** Anju K. Ojha, Pawan K. Dhar

## Abstract

This study explores the intricate world of protein-protein interaction (PPI) networks and the critical role of hub proteins within them. PPIs, mediated by various biomolecular forces, orchestrate diverse cellular processes through transient or stable interactions. The study delves into the unique ability of hubs to connect with numerous partners within the network. This research investigates the network properties and focuses on hub-centric questions. This study explores information-carrying capacity, power laws in protein circuits, connectivity profiles of hub proteins, and network expansion or shrinkage during evolution. The study validates the power law for connectivity, identifies a marginal network compression in evolution, unveils hub connectivity patterns, and encourages the exploration of the correlation between protein structure, connectivity, and function. These findings offer valuable insights into network design principles and the critical role of hub proteins in cellular function and evolution.

## INTRODUCTION

Proteins are macromolecules composed of amino acid chains that fold into unique three-dimensional structures. These structures dictate a protein’s specific function, often involving binding to other molecules, particularly other proteins. Protein-protein interactions (PPIs) are specific linkages mediated by various forces, including hydrogen bonding, electrostatic interactions, and hydrophobic effects. These interactions create transient or stable protein complexes with distinct biological functions. The tightly bound, long-lasting interactions often form the scaffolding for permanent protein complexes, essential for cellular structures like ribosomes and channels. Similarly, transient interactions serve as dynamic switches, triggering signal transduction pathways or regulating enzymatic activity.

Protein-protein interaction networks are graphical representations in which proteins are depicted as nodes and interactions between them as edges. The strength and type of interaction can be further incorporated as edge attributes, enriching the network with additional information. PPI networks are dynamic representations teeming with biological information. The network topology, the arrangement of nodes and edges, provides valuable insights into cellular organization. Highly connected nodes, called hubs, represent proteins with critical roles in coordinating cellular processes. Identifying and analyzing these hubs can lead to a deeper understanding of essential cellular pathways.

Deeply embedded in the intricate network of biomolecular interactions, are hub proteins. These unique molecules act as connectors, interacting with many partners within the cellular PPI network. The defining feature of a hub protein is its ability to bind to many other proteins. Hub proteins occupy central positions within the PPI network, acting as bridges between different functional modules. Studies suggest that hub proteins are often more critical for survival than their less-connected counterparts. The high connectivity of hub proteins makes them ideal candidates for scaffolding and Orchestration, signal integration and transduction and maintaining network integrity.

This study aimed to move from overall network properties to hub-centric transactions, hoping to uncover regularities to identify new network design principles. Questions addressed in the current study were: given the increased biological complexity during evolution, has the information-carrying capacity per network pixel also increased? Do protein circuits exhibit power law? What is the connectivity profile among hub proteins in the PPI networks? Have networks expanded or shrunk in evolution? If yes, how do expansion/compression hotspots look? The current study looks into some aspects of network biology and tries to find future research hotspots that may lead to the uncovering of organizing principles.

## MATERIAL AND METHODS

Six organisms, H. pylori, E. coli, S. cerevisiae (Yeast), C. elegans (Worm), D. melanogaster (Fly) & H. sapiens (Human) were selected due to their rich annotation and interaction data. The publicly available protein-protein interaction databases (DIP, BIND, IntAct, Reactome, HPRD & MINT DBs were manually used in this study. Ensembl was used to find ortho proteins. Exact sequence homology was utilized to reconcile unique accession identifiers across databases, resulting in a meta-dataset from which a non-redundant set of proteins and interactions was extracted.

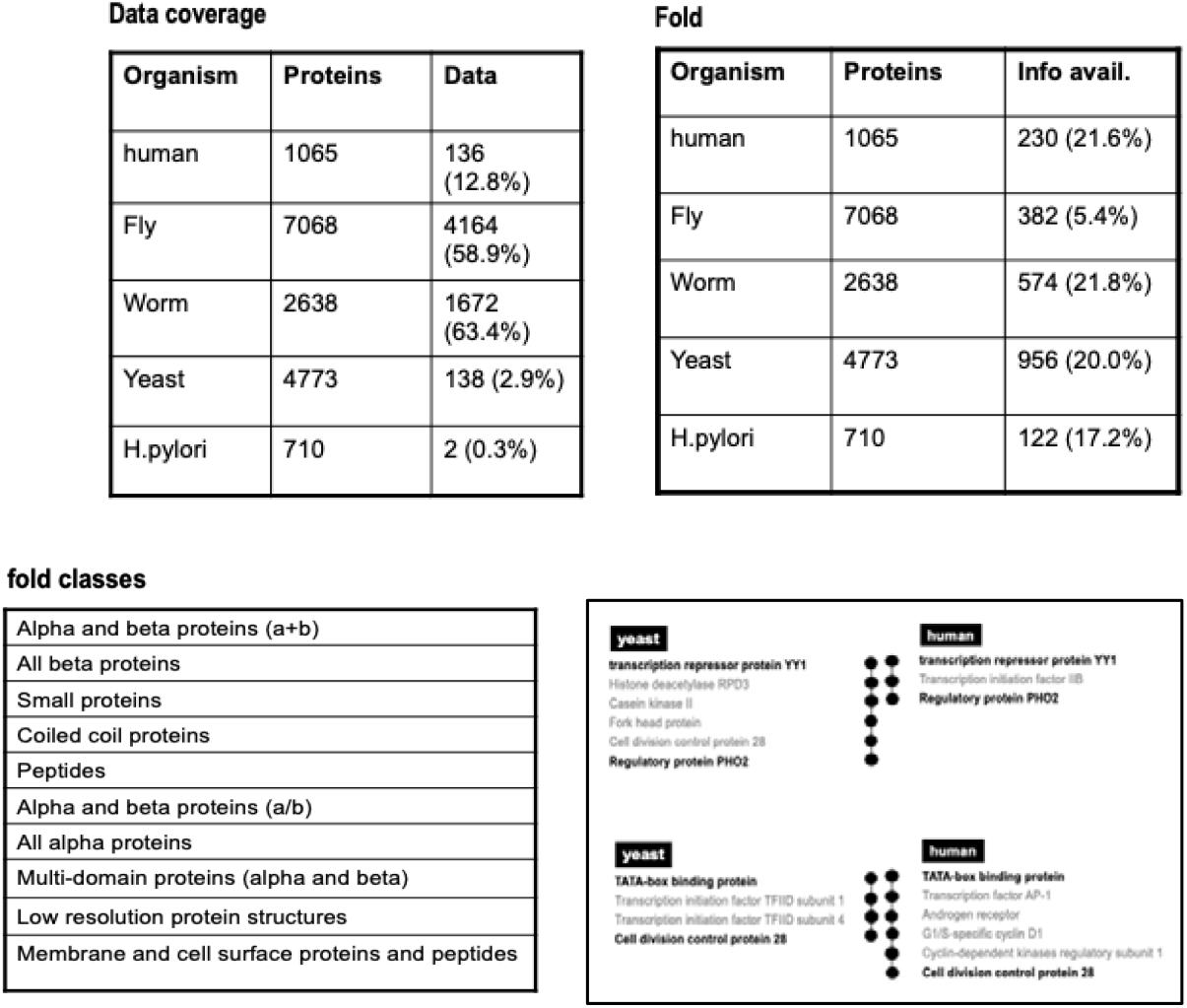

The GO functions considered in the present study were: antioxidant activity, binding, catalytic activity, chaperone regulator activity, enzyme regulator activity, molecular function unknown, motor activity, nutrient reservoir activity, protein tag, signal transducer activity, structural molecule activity, transcription regulator activity, translation regulator activity, transporter activity and triplet codon-amino acid adaptor activity

Pairwise orthologs were then clustered into multi-species groups. Identical proteins with different annotation names were identified through global sequence comparison and assigned a unique ID. A global sequence similarity score of more than 60 was used to group two different proteins under the same name. The connectivity score of proteins in other organisms was tabulated. Fold change definition for connectivity (kfc = k/hki) was selected, along with appropriate cut-offs to identify hubs in different PPI networks. Prokaryotic hubs were identified with kfc ≥ 2 (cutoff, P < 0.03), while for eukaryotes, kfc ≥ 10 (with P < 0.001). Stringent hub criteria were applied to orthologs, and only those meeting these criteria in at least one species were selected.

Three classes of hubs were identified based on their connectivity trends across species: ‘getting rich’ hubs, ‘getting poor’ hubs, and ‘flexible’ hubs with non-uniform connectivity trends. Gene Ontology (GO) annotations for each protein were obtained from source databases (BIND, SGD, Flybase, Wormbase, HPRD), enriched into ten top-level molecular functions.

The study also aimed to determine if hub proteins retained the same partners across species. A statistically significant similarity (e-value < 0.001) indicated partner retention, while lack thereof was considered a partner change. To calculate an average score, individual connectivity computed and normalized the total number of evolutionarily conserved interactions. Statistical significance of the identified hub classes was assessed using F-test.

The shortest protein-protein interaction pathways were identified and compared. To determine expansion/contraction, terminal nodes were kept consistent. Terminal nodes with different IDs were identified through sequence comparison. Next, the shortest path between any two terminal nodes was determined. The overall strategy was to identify a protein’s hubness, find its connectivity patterns across phylogeny, and link the connectivity data with structure and function data in search of regularities.

## RESULTS

Given the increased biological complexity during evolution, we asked if the information-carrying capacity per network pixel also increased proportionately. To answer this question, we compared the connectivity score of each protein and metabolite of model organisms. As is evident from Table 1, there was a proportional increase in the connectivity score of metabolites from H. Pylori to H. Sapiens. The growth is observed as an average link per node and in the range of links. However, the same was not found for proteins, perhaps due to low data density. Next, we asked: Do protein circuits exhibit power law? Figure 1 shows an interesting trend. A power law (preferential attachment model) was observed in most organisms. As is evident from Figure 1, the log-log plot shows protein molecules following an overall linear distribution. However, in S.cerevisiae protein molecules seem to be falling off the axis.

**Table 1:** An evolutionary trend of metabolic and protein-protein interactions.

|  | Number of nodes | Average no. of links / node | range of links |
| --- | --- | --- | --- |
| Helicobacter pylori | 576 | 5.26 | 1-119 |
| Saccharomyces cerevisiae | 969 | 5.49 | 1-251 |
| Caenorhabditis elegans | 898 | 5.32 | 1-225 |
| Drosophila melanogaster | 937 | 5.22 | 1-220 |
| Arabidopsis | 1101 | 5.32 | 1-268 |
| Mouse | 1219 | 5.78 | 1-335 |
| Homo sapiens | 1339 | 6.08 | 1-370 |

|  | Number of nodes | Average no. of links / node | range of links |
| --- | --- | --- | --- |
| Helicobacter pylori | 710 | 2.0 | 1-54 |
| Saccharomyces cerevisiae | 4773 | 3.24 | 1-187 |
| Caenorhabditis elegans | 2638 | 1.52 | 1-187 |
| Drosophila melanogaster | 7068 | 2.97 | 1-178 |
| Homo sapiens | 1065 | 1.29 | 1-34 |

**Table 2:** Distribution of folds and function among clustered regions found in D.melanogaster.

| Fold | Region I | Region II | Region III |
| --- | --- | --- | --- |
| All alpha proteins | 0 | 3 | 50 |
| All beta proteins | 0 | 2 | 34 |
| a+b | 1 | 5 | 81 |
| a/b | 2 | 1 | 49 |
| Coiled coil proteins | 0 | 0 | 6 |
| Membrane and cell surface proteins | 0 | 0 | 2 |
| Multi-domain proteins | 0 | 0 | 5 |
| Peptides | 0 | 0 | 3 |
| Small proteins | 0 | 0 | 21 |

| Function | Region I | Region II | Region III |
| --- | --- | --- | --- |
| antioxidant activity | 0 | 0 | 11 |
| binding | 25 | 26 | 1687 |
| catalytic activity | 12 | 14 | 1120 |
| enzyme regulator activity | 0 | 3 | 111 |
| molecular function unknown | 1 | 1 | 89 |
| motor activity | 0 | 0 | 25 |
| nutrient reservoir activity | 1 | 0 | 4 |
| signal transducer activity | 6 | 3 | 240 |
| structural molecule activity | 7 | 5 | 303 |
| transcription regulator activity | 6 | 7 | 380 |
| translation regulator activity | 0 | 1 | 46 |
| transporter activity | 1 | 2 | 302 |

**Figure 1:**
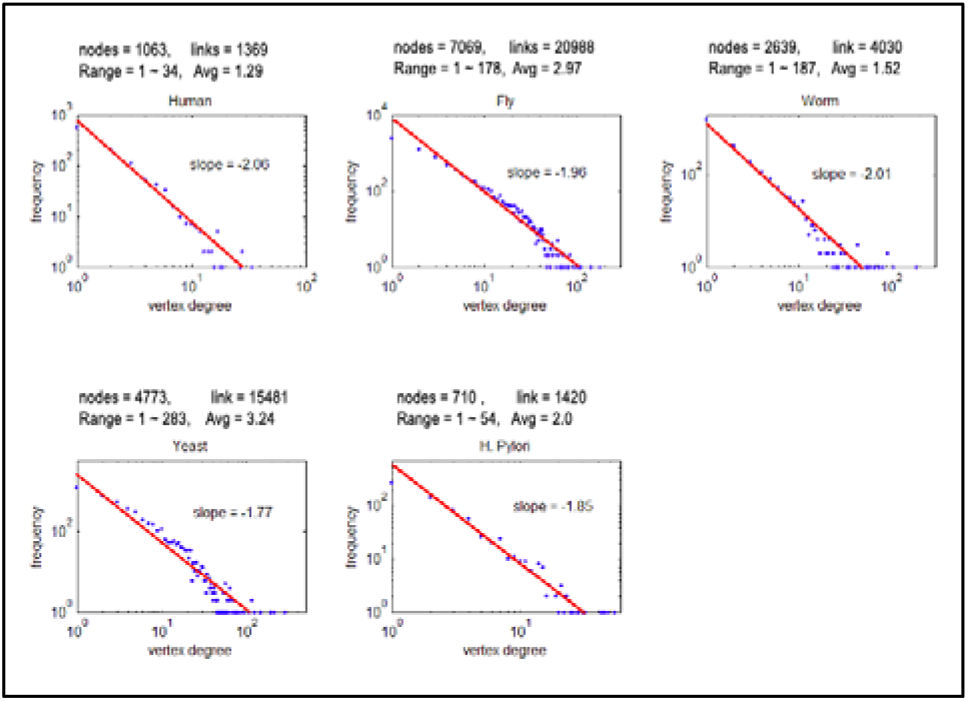
The log-log plot of protein connectivity across various organisms

**Figure 1:**
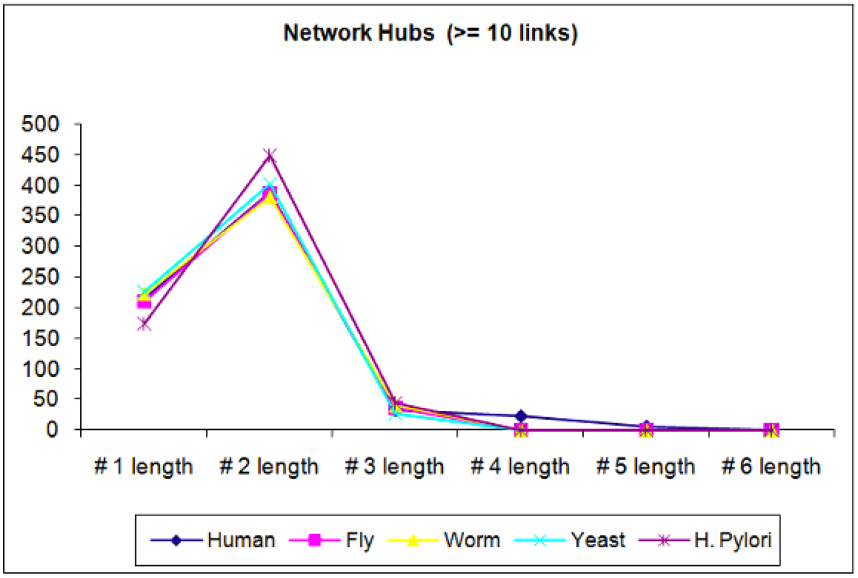
The hub-hub protein connectivity pattern

To address the question of inter-hub communication, we plotted the number of linker nodes among all hub proteins on the x-axis and their frequency on the y-axis. It was remarkable to observe an amazing consistency among all the organisms studied. In most cases, hub proteins were connected through two protein molecules (linker). Next in the decreasing order of frequency was one linker protein occurring between any two hub proteins. Following, there was a long tail that also showed amazing similarity among organisms studied (Fig 2). The figure strongly points to a design principle consistently followed by the protein-protein interaction networks. It was interesting to find that most of the hubs were connected by two linker proteins in between. Next in the line was single connectivity between hub proteins. The rest (above three linker proteins) was a long tail, remarkably conserved among organisms.

**Figure 2:**
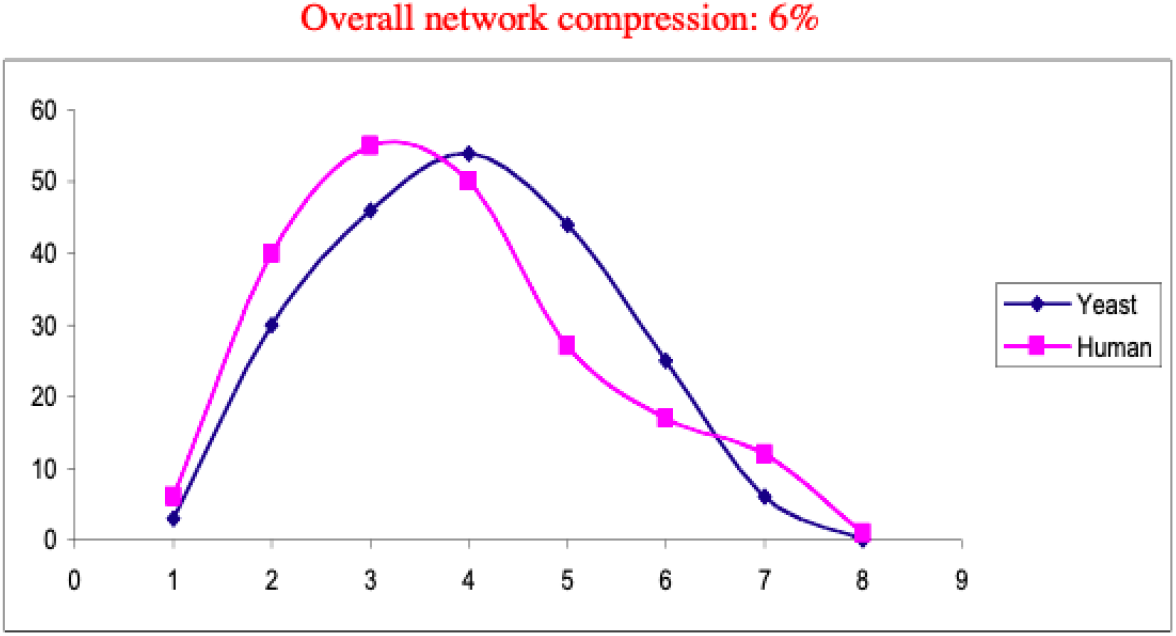
Data from 35 shortest pathways with

**Figure 4:**
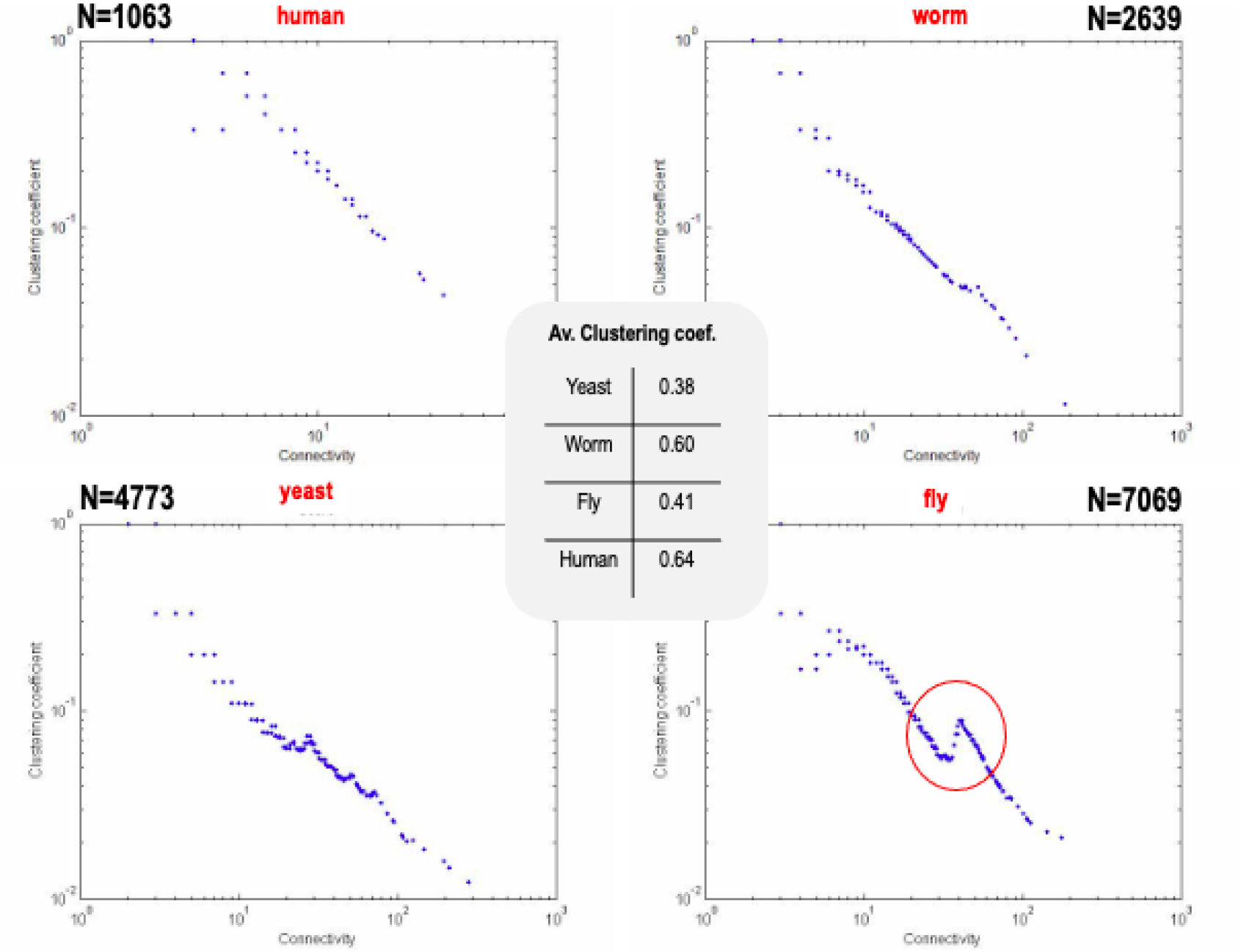
Clustering patterns found in PPI networks

Next, we asked: Have networks expanded or shrunk in evolution? If yes, how do expansion/compression hotspots look?

Data suggested that the PPI network diameter may have shrunk with evolutionary complexity (Fig 3). This trend does not take into account functionalization of network expansion/compression. The reasons for this observation are unclear. More studies need to be performed to figure out evolutionary reasons.

Next, we asked: Do protein circuits exhibit unique clustering patterns?

The current data (n=1063 to 7068) revealed no clustering patterns in humans and worms. However, in the yeast, rudimentary clustering peaks started showing up, which was significantly observed in the flies. Two subpopulations were demarcated and were joined by a narrow bridge.

Questions that arose from clustering patterns were: Do these subpopulations have a structural basis? Do they represent special functional zones? A deeper analysis indicated that Individual protein folds and functional patterns did not explain the phase transition in D.melanogaster PPI networks.

## DISCUSSION

The organizing principles of biology provide a framework for understanding the natural world. By appreciating these principles, we understand how living systems function at different levels, from the molecular to the ecological. As we explore the intricate dance of life, these organizing principles will continue to guide our scientific inquiry, helping us unlock the mysteries of life’s extraordinary complexity.

The question is: Can one go beyond Mendel’s inheritance laws and discover new laws or organizing principles of biological complexity? If yes, what’s the right approach to find regularities and weave them into convincing stories? To address these and more questions, we considered protein-protein interaction networks and studied them at the connectivity level and at the structural and functional levels.

PPI networks provide a robust framework for understanding standard homeostatic mechanisms and addressing disease conditions. Several experimental techniques have been instrumental in constructing PPI networks used in the present study, e.g., Affinity chromatography, Competition binding, Gel filtration chromatography, Genetic screening, Immunoblotting, Immunofluorescence, Immunolocalization, Immunoprecipitation, In vitro binding, NMR, Surface plasmon resonance, Two hybrid test and, X-ray diffraction. Beyond experimental approaches, computational methods are crucial in PPI network construction. Homology-based methods infer interactions based on sequence or structural similarities to proteins with known interactions. Text mining algorithms browse scientific literature to identify protein interactions. Integrating data from various sources allows for more comprehensive and robust PPI networks. However, each technique has its limitations, leading to false positives and negatives. However, enlarging the data size can minimize variations and rely better on the outcomes.

It is worth noting that the data used in the present study was a partial representation of the system, taking into consideration the data that came up in the form of superimposition of culture conditions, strains, metabolic state, and time kinetics. The PPI data considered here was studied in relation to the information-carrying capacity per node, protein resource allocation to support various functions, power law, and connectivity-structure-function correlations.

We hypothesized that as systems become more complex, from prokaryotes to eukaryotes, information carrying capacity per node will increase, and connectivity patterns will show some preferential correlation. The data showed an evolution-correlated increase in connectivity scores for metabolic networks but not PPI networks.

Next, we investigated the properties of PPI networks to follow power law. Protein connectivities were plotted on the log-log curve. Most of the organisms seem to follow the power law. However, in the yeast the molecules seem to fall off after traveling half the distance. The reason for this needs to be clarified. More data is required to validate this observation.

Next, we wanted to see the shortest path between hubs, taking into consideration a single linker protein, two linker proteins, three linkers, and so on. The data shows a remarkable correlation that is evolutionarily conserved. Most organisms show one linker protein connecting the hubs, followed by two linker proteins, and so on. This noteworthy design principle has been evolutionarily conserved and points to the need for maintaining robust communication among hubs.

Further, the theme of network expansion/compression was explored. We worked on the assumption that, with evolution, networks would have expanded to handle more complex tasks. The data shows an overall network compression of 6% from yeast to humans. In the future, this observation needs to be examined in detail by including functional domains of the PPI networks.

Finally, we asked if PPI networks exhibited unique clustering patterns. The PPI networks were plotted using the connectivity vs. clustering coefficient, and a distinct pattern was observed in the yeast PPI network. On closer examination of three regions in D. Melanogaster, a foundational correlation with structure and/or function was not observed. The clustering coefficient also used a nonuniform evolutionary correlation from E. Coli to H.sapiens. From the data available, individual protein folds and functional patterns do not explain the “phase transition” observed in the fly PPI networks.

Overall, the current study points to design principles that merit deeper exploration. The study is preliminary and based on manually acquired data. In the future, more work is required to include a larger dataset, review this observation, and ask questions across three levels: structure, connectivity, and function. This is an initial study that shows interesting correlations. In the future, more work is required to validate these observations with more data and explore the structural basis of connectivity and functional attributes.

